# Long-range neural coherence encodes stimulus information in primate visual cortex

**DOI:** 10.1101/2020.06.22.164269

**Authors:** Mojtaba Kermani, Elizabeth Zavitz, Brian Oakley, Nicholas S.C. Price, Maureen A. Hagan, Yan T. Wong

**Affiliations:** Department of Physiology, Biomedicine Discovery Institute, Monash University, Clayton, VIC 3800, Australia; Department of Electrical and Computer Systems Engineering, Monash University, Clayton, VIC 3800, Australia

**Keywords:** Marmoset, V1, spike-LFP coherence, synchronization, mutual information

## Abstract

In the primary visual cortex, neurons with similar receptive field properties are bound together through widespread networks of horizontal connections that span orientation columns. How connectivity across the cortical surface relates to stimulus information is not fully understood. We recorded spiking activity and the local field potential (LFP) from the primary visual cortex of marmoset monkeys and examined how connectivity between distant orientation columns affect the encoding of visual orientation.

Regardless of their spatial separation, recording sites with similar orientation preferences have higher coherence between spiking activity and the local field potential than sites with different preferred orientation. Using information theoretic methods, we measured the amount of stimulus information that is shared between pairs of sites. More stimulus information can be decoded from pairs with the same preferred stimulus orientation than the pairs with a different preferred orientation, and the amount of information is significantly correlated with the magnitude of beta-band spike-LFP coherence. These effects remained after controlling for firing rate differences.

Our results thus show that spike-LFP synchronization in the beta-band is associated with the encoding of stimulus information within the primary visual cortex of marmoset monkeys.

**Significance Statement:** A fundamental step in processing images in the visual cortex is coordinating the neural activity across distributed populations of neurons. Here, we demonstrate that populations of neurons in the primary visual cortex of marmoset monkeys with the same stimulus orientation preference temporally coordinate their activity patterns when presented with a visual stimulus. We find maximum synchronization in the beta range depends on the similarity of orientation preference at each pair of the neural population.

## Introduction

The columnar hypothesis proposes that sensory cortices are comprised of highly connected modular columns of neurons. In the visual cortex, these columns are formed by clusters of neurons with similar receptive field properties such as orientation preference (Mountcastle, 1997). A fundamental step in analyzing a visual object is combining different features of the object such as orientation which are encoded in spatially separate functional columns.

To combine encoded features, it has been proposed that neurons use intricate networks of long-range horizontal connections which often preferentially connect clusters of neurons that have similar orientation preference (Gilbert and Wiesel, 1989; Kisvarday et al., 1989; Kisvárday and Eysel, 1992; Bosking et al., 1997). Such functional specificity may allow single neurons to combine information from a larger area in the visual field (i.e. neighboring orientation columns) than that covered by their individual receptive fields (Burkhalter, 1989; Schwarz and Bolz, 1991; Malach et al., 1993, 1993; Bosking et al., 1997, Anon, 2013; Liang et al., 2017).

Cross column processing requires temporal structure similarity between columns (Fries, 2005, 2015; Womelsdorf et al., 2007; Fries et al., 2008) and neural synchrony has been theorized as a major contributor for the precise timing required for neural communications (Buzsáki and Draguhn, 2004; Fries, 2005, 2015; Dean et al., 2012; Hawellek et al., 2016; Wong et al., 2016). Earlier studies of the visual cortex of cats have supported this idea by showing that spatially-separated, orientation-selective neurons with the same stimulus preference synchronize the pattern of their spiking activities when they are stimulated with a visual stimulus (Gray et al., 1989). Spike threshold have been shown to be closely correlated with fluctuations in membrane potentials of cortical neurons in a frequency dependent manner, where spiking activity exhibit a higher correlation with high-frequency components of membrane potential than the mean membrane potential (Azouz and Gray, 2003). Therefore, the local field potential (LFP), which is combined electrical activity in the extracellular medium (Mitzdorf, 1985; Einevoll et al., 2013) may play a key role on neural synchronization through modulating the excitability of neurons (Buzsáki and Draguhn, 2004; Buzsaki and Schomburg, 2015; Fries, 2015).

The relationship between spikes and the LFP in the visual cortex has been quantified using spike-LFP coherence (SFC), which is the degree of correlation between spikes and different frequency bands of the LFP. The idea of communication through coherence (CTC) between spikes and the LFP has been studied in several studies. SFC in visual area V4 is predictive of monkey’s reaction times for detecting stimulus changes (Womelsdorf et al., 2006). Perception of stimuli during interocular rivalry can also be tracked by the degree of SFC in primary visual cortex (V1) neurons (Fries et al., 1997). Visual attention increases SFC between neurons within area V4 (Fries et al., 2008) and between area V4 and prefrontal cortex neurons (Gregoriou et al., 2009). Layers of the visual cortex (V1, V2 and V4) exhibit SFC in different frequency bands (Buffalo et al., 2011; Lashgari et al., 2012). Although these studies have proposed synchronization between visual neurons as the way of establishing channels for neural communication which can be tracked by SFC, how SFC across the cortical surface relates to stimulus information is not fully understood.

We propose that in the visual cortex, communication between functional columns is facilitated when spikes in one column are synchronized with the LFP in another column. Therefore, higher SFC magnitude is expected to be extracted from pairs of columns with the same stimulus orientation preference as they are more interconnected than pairs of columns with different orientation preferences.

We simultaneously recorded visually-evoked spikes and LFPs from V1 of anesthetized marmoset monkeys. We used spike-LFP coherence and information-theoretic methods to quantify the degree of synchronization within and between orientation columns. Sites with similar orientation preferences had higher spike-LFP coherence than sites at the same spatial separation with different preferred orientations. We measured the amount of stimulus information that is shared between pairs of sites and found that pairs with the same preferred stimulus orientation hold more stimulus information than the pairs with different preferred stimulus orientation. The amount of shared information between pairs is also significantly correlated with beta-band spike-LFP coherence (22 Hz). This suggests that CTC is a plausible mechanism for inter-column information processing.

## Methods

### Material and methods

Four adult marmoset monkeys (3 male, 1 female; Callithrix jacchus) were used in this study. All experimental procedures were approved by the Monash University Animal Ethics committee and were conducted in accordance with the Australian Code of Practice for the Care and Use of Animals for Scientific Purposes. After the initial induction of anesthesia using Alfaxalone (8mg/kg, IM), a tracheotomy, vein cannulation and craniotomy were performed (Zavitz et al., 2016). After paralyzing the skeletal muscles by using an intravenous bolus of paralytic (Pancuronium Bromide; 4 mg), the animal was artificially ventilated with a mixture of nitrous oxide and oxygen (7:3). Anesthesia was maintained by the intravenous administration of Sufentanil.

Pupillary dilation was achieved with the topical application of atropine (1%, 1 drop/eye) and phenylephrine hydrochloride (10%; 1 drop/eye). Eyes were fitted with a pair of rigid gas-permeable contact lenses to prevent corneal drying and appropriate corrective spectacle lenses were used to bring a display at 1 m into focus.

A craniotomy opening was made over V1. Following a durotomy, a 96-channel “Utah” array (Blackrock Microsystems) with 1.5 mm electrodes was implanted in each animal using a pneumatic insertion tool. The location of the array was chosen based on gross anatomical landmarks and was verified histologically and based on the characteristics of neurons’ receptive fields. The ipsilateral eye was occluded to achieve monocular vision.

### Visual stimuli

Visual stimuli were generated by Psychtoolbox running on MATLAB and presented on an LCD monitor with linear gamma (Display++, Cambridge Research Systems, UK; 700 mm display width; 1920 × 1080 pixels refreshed at 120 Hz) positioned at a viewing distance of 1 m. The stimuli consisted of drifting full contrast sine wave gratings in a circular aperture with 12 equally spaced directions (six orientations), presented for 500 ms followed by 500 ms grey screen after each trial.

Neurons were characterized based on receptive field location (using flashed squares, 0.25-1 degree), spatial frequency (between 0.2 and 2 cycles per degree) and temporal frequency (between 1 and 10 Hz) tuning. The spatial and temporal frequency that best evoked responses across the array were chosen and run across stimulus directions where each direction was randomly repeated 75 times in each block of trials (Fig. 1a). The size of the stimuli was selected to cover all neurons’ receptive fields across the array and ranged from 4.5 to 6 degrees of visual angle.

**Fig. 1.**
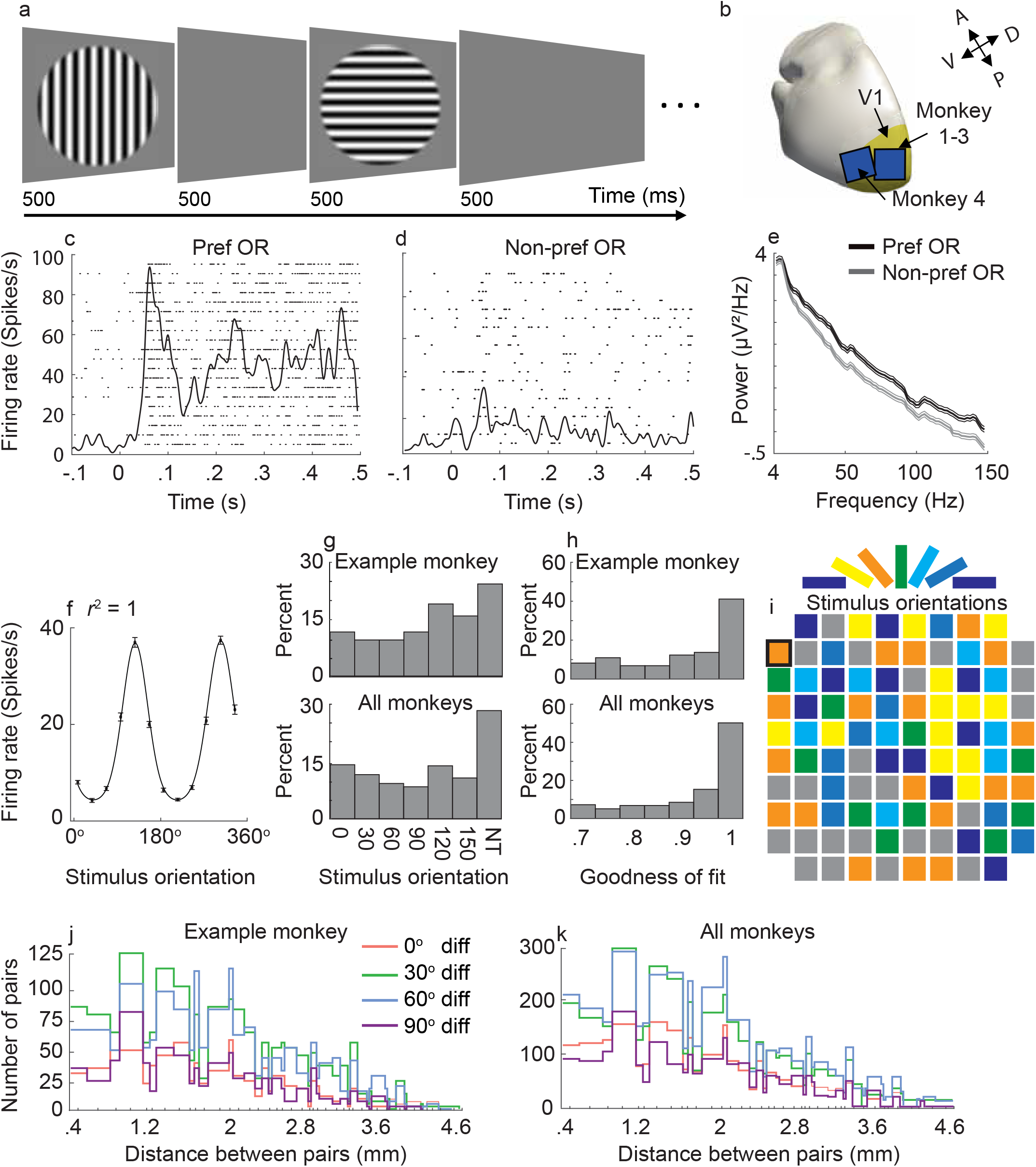
Stimulus design and neural responses of V1 neurons. **a**, A random sequence of oriented gratings was presented. Each grating lasted 500 ms followed by a 500 ms grey screen, and each direction was presented 75 times. **b**, Locations of electrode implantations. Three electrodes covered the entire V1 while the forth covered V1 and V2. **c,d**, Responses of an example site to its preferred and non-preferred orientation. **e**, Spectral power distribution of an LFP site to the preferred and non-preferred stimulus. The thin lines represent standard error of the mean. **f**, Orientation tuning for the example unit. The site has the maximum goodness of fit (r^2^ = 1). **g**, Distribution of orientation preference for the example site and population. NT: non-tuned sites. **h**, Distribution of the goodness of fit for the example site and population. Only sites with significant orientation tuning are included. **i**, Orientation map across the Utah array for an example animal. Sites without orientation preference are marked in gray. The site with a black margin represents the example site which is used in c-f. **j,k** Number of paired sites in each preferred-orientationdifference group for each distance. The orientation of the stimulus is color-coded and the nontuned sites are marked grey

### Electrophysiology

The raw voltage signal was recorded with a Cerebus multichannel data acquisition system (Blackrock Microsystems) and was filtered at 0.1 – 4000 Hz with a sampling rate of 30 kHz. To extract spikes and LFP from each electrode, the recorded signal was band-pass filtered in the 4 - 100 Hz range for LFP and in the 300 – 4000 Hz range for the spiking activity. Spike density functions were derived for each site by binning the spike trains in 50 ms bins and then calculating a trial-averaged rate smoothed by a Gaussian kernel with 50 ms width for better visualization. The PSTHs at each of the orientation directions were plotted to visualize the responsiveness of sites to the stimuli.

In one monkey (M 1), we extracted single units using Wave_clus toolbox running on MATLAB (Quiroga et al., 2004), which is a semi-automatic spike detection and sorting toolbox using wavelets and super-paramagnetic clustering. Spikes were detected with a pre-defined amplitude threshold (3*standard deviation of baseline noise) and were clustered based on their wavelet coefficients. We found that the peak coherence computed for sorted spikes and multiunit threshold crossings were not significantly different (p = 0.038, Wilcoxon rank-sum test). Therefore, for that animal and the other three animals, we only used multiunit data for all further analyses (Trautmann et al., 2019).

### Orientation tuning

For each test direction, we calculated the mean firing rate throughout the stimulus presentation (500 ms) and plotted them together to yield the tuning curve for each recording site. Orientation sensitivity was estimated by fitting a *Von Mises* function described by Equation 1: (Swindale et al., 2003)

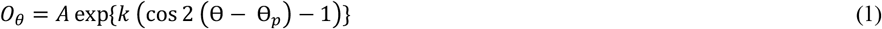

Where, O is the response of the model for the stimulus orientation Θ, A and k are the maximum height and width of the orientation tuning curve and Θ_p_ is the preferred orientation.

Only sites where the fit had an *r*^2^ value larger than 0.7 and showed a significantly better fit than a straight-line function (*p*-value < 0.05, *F* test) were identified as orientation-selective and included for further analysis.

### Spike-LFP coherence

The coherency spectrum between spiking activity and LFP was measured using Equation 2 (Mitra and Pesaran, 1999):

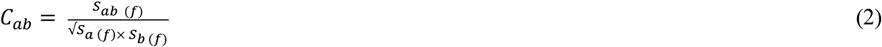

Where, S_a_, S_b_ and S_ab_ are spectra of spikes, the LFP and their cross-spectral densities. The coherence magnitude ranges from zero to one where a magnitude of one denotes maximum correspondence of the two. We used the multitaper method (Thomson, 1982), implemented in Chronux 2.0 (Bokil et al., 2010) with a 500 ms analysis window and 6 Hz smoothing aligned to the onset of the stimulus.

### Information-theoretic analysis

We quantified how much information about the stimulus is shared between sites by using the information theoretic analysis method (Shannon, 1948). We calculated the mutual information (MI) using Equation. (3):

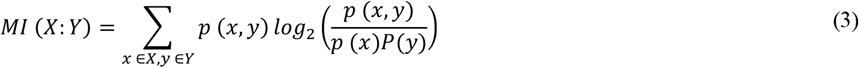

Where, X and Y are spike counts for site 1 and 2, p(X) and P(Y) are the probabilities of observing the response in the site 1 and 2 evoked by the stimulus during 50 - 350ms stimulus window. Information theoretic measures suffer from sampling or stimulus-response bias (Panzeri et al., 2007). To achieve an unbiased estimation of information, we performed bootstrap-based bias-correction method (Panzeri et al., 2007). We also performed significance testing on estimated MI using Monte Carlo analysis (Timme and Lapish, 2018) and only pairs with p < 0.05 were included for further analysis.

### Spike-Count Correlation Measurements

Spike-count correlations (*r._sc_*) were calculated as the Pearson correlation between spike counts of trial-averaged responses to 12 directions for each pairings, counted over the entire duration of the stimulus.

### LFP phase difference analysis

To quantify the LFP phase difference between pairs, we used a wavelet-based filter to isolate the 22 Hz frequency components of the LFP. Then we applied the fast Fourier transform to estimate the phase of this complex signal throughout stimulus presentation and quantified the phase difference for pairs with distance from 400 to 1260 μm.

### Statistical analysis

Non-parametric permutation testing was used to assess the statistical significance of the magnitude of the spike-LFP coherence in each frequency band. In each group, the order of trials in LFP sites was shuffled 1000 times to create a null distribution. The initial significance was assessed by comparing the actual spike-LFP coherency (SFC) magnitude values with that of the shuffled dataset. A cluster-based correction was used for multiple comparisons (Maris and Oostenveld, 2007). All circular statistics were performed using custom-made MATLAB scripts.

## Results

Spiking activity and local LFP potentials (LFPs) in response to visually presented oriented gratings (**Fig 1a**) were simultaneously recorded from 96-electrode Utah arrays implanted in the primary visual cortex (V1) of four anaesthetized marmoset monkeys (**Fig. 1b**). Neurons showed a brief (50-100 ms) transient surge in their firing rate, which was followed by a sustained response for the duration of the stimulus presentation (**Fig 1c,d** for preferred and non-preferred orientations, respectively). The stimulus orientation resulting in a maximum spiking response was selected as the preferred orientation. An example neural response to the preferred orientation for twenty trials are represented as a raster plot in Figure 1c for the preferred orientation and Figure 1d for its perpendicular (non-preferred) orientation. LFP spectral power was also modulated by changes in stimulus orientation (**Fig. 1e**; same site as c, d and f).

We recorded from 280 visually responsive sites (< 0.05, t-test, pre- vs post-stimulus onset firing rates) with 221 (79%) of these showing significant orientation tuning (61, 53, 66 and 41 sites from animals 1-4, respectively). **Figure 1f** illustrates an example of a tuning curve from one recording site with significant (r^2^= 1, p = 1.8e-09, F-test) orientation tuning. Only sites with significant orientation tuning were included for further analysis. The proportion of tuned sites for each stimulus orientation was different for each animal, however, there was a minimum of seven tuned sites for each stimulus orientation in each animal. Pooling data across four animals, we found even representation across all orientations which made our dataset valid for statistical analysis. (**Fig. 1g**).

To inspect the quality of our recordings, we measured the distribution of the goodness of fit for sites with significant orientation tuning (**Fig. 1h**). The tuning was robust with 111 sites showed the maximum orientation tuning (r^2^ = 1). Tuned sites were quasi-randomly distributed across the Utah array (**Fig. 1i** for an example monkey, M1).

We then paired recording sites and compared their stimulus orientation preference. We recorded 11,225 pairs of orientation-tuned electrodes across all monkeys (3,012, 2,292, 3,498, and 2,423 for monkeys 1 to 4, respectively). The difference in preferred orientation for each electrode pair was determined (0, 30, 60 or 90-degree preferred orientation difference) as well as the cortical distance between each electrode pair. Distances between electrodes across the array ranged from 0.4 mm (adjacent electrodes) to 5.1 mm with large numbers of pairs for all preferred orientation differences across animals (**Fig. 1j**, example monkey; **Fig. 1k**, all monkeys).

### Spike-LFP pairs with the same preferred stimulus orientation exhibit stronger coherency

Neural synchronization is a proposed mechanism of enhancing communication between remote neuronal populations. We hypothesized that columns with the same orientation preference should, therefore, exhibit robust synchronization and, consequently, highest communication as spikes sent by one cluster arrive in the other neural cluster during the time window of their maximal sensitivity. To test whether the strength of synchronization between clusters of neurons was dependent on stimulus orientation preference (**Fig 2a**), we measured synchronization across cortical columns, by calculating spike-LFP coherence between pairs of electrodes using the spiking activity on one electrode (spike sites) and the LFP on the other electrode (LFP sites), averaged across all stimuli and all trials.

**Fig. 2.**
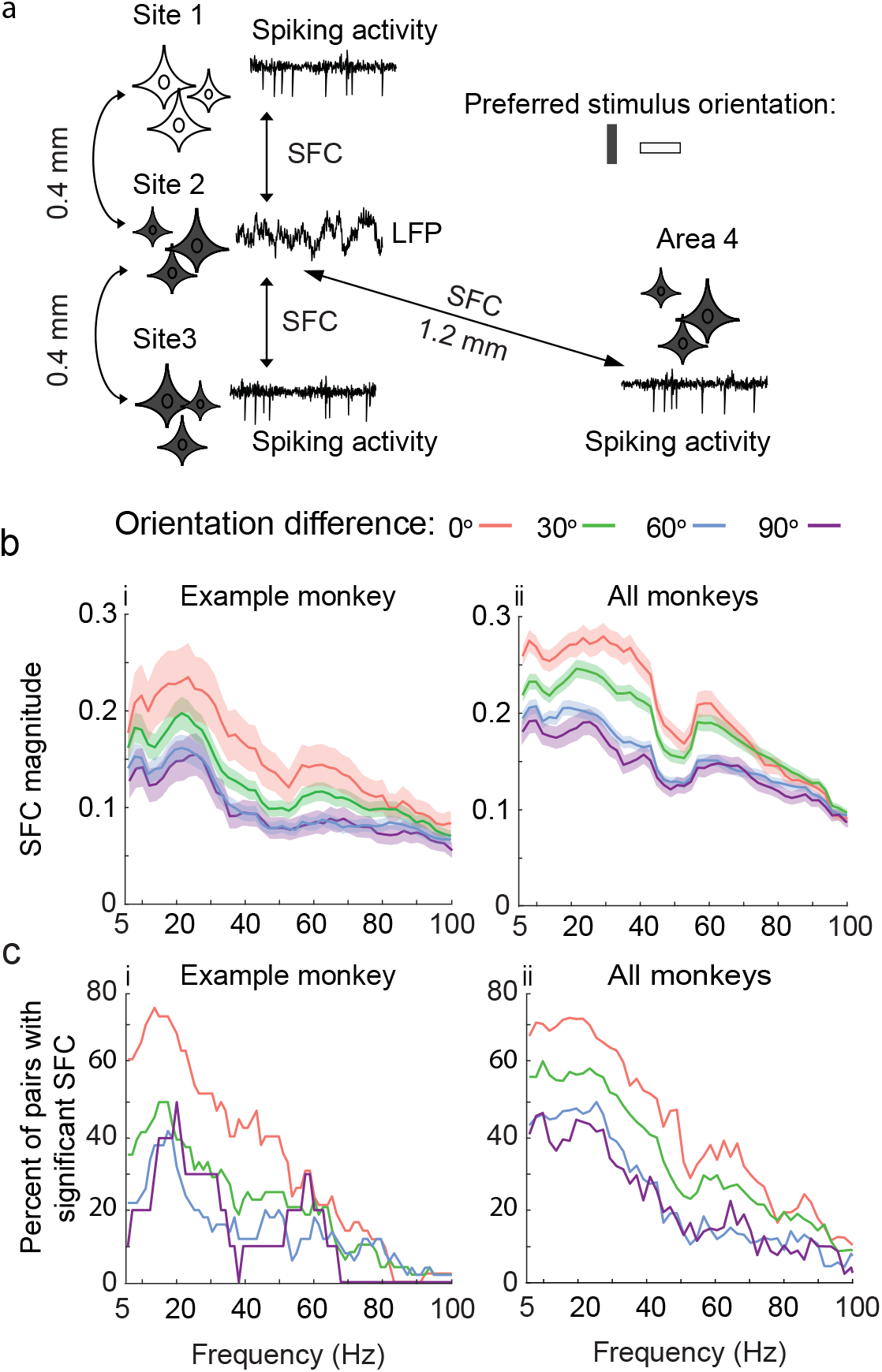
Spike-LFP coherency of neighboring sites. **a**, Conceptual diagram outlying functional connectivity of neighboring pairs with similar tuning. Distance between sites 1-3 are equal, but the strength of functional connectivity, as the magnitude of spike-LFP coherence, between sites 2 and 3 is higher than between sites 1 and 2 since the first pair has the similar preferred stimulus. Remote sites (site 4) with the same stimulus preference are also functionally connected together. **b_i,ii_**, Spike-field coherence magnitude as a function of frequency between sites with a 400-micrometer distance in the example monkey and the population. The coherency between pairs with the same preferred orientation is higher than that of those with different stimulus preference. **c_i,ii_**, The percentage of pairs that reached statistical significance at each frequency bin for the example monkey and population. A higher percentage of reaching the statistical significance was observed when pairs had the same preferred orientation preference. Pairs with the same preferred stimulus orientation show a higher percentage of significant SFC.

Spike-LFP pairs were grouped based on the difference in preferred orientation between the spiking activities on each electrode (0, 30, 60 or 90-degree). We labeled the orientation preference of a channel using the tuning curve from spiking on that channel and not the preference of the LFPs themselves as similar with previous studies (Lashgari et al., 2012), we found the same orientation tuning for LFP and spiking activity in the gamma range (40-100Hz) in 88.6 % of sites.

We first quantified the spike-LFP coherence (SFC) of neighboring electrode sites (400 μm distance) with the same orientation preference. Spike-LFP coherence peaked in the beta-band in all pairing groups (20-30 Hz **Fig. 2bi**, example monkey, Monkey 2, n = 358 electrode pairs; **Fig 2bii** all monkeys, n = 1,091 electrode pairs). It should be noted that electrode pairs are double-counted (e.g. spike-LFP and LFP-spike).

SFC magnitude decreased when LFP and spike sites had different preferred stimulus orientation. The amount of attenuation depended on the difference in preferred orientation so that pairs with the same preferred orientation (0-degree difference) showed the highest SFC, followed by pairs with 30-degree preferred orientation difference, and then pairs with 60- and 90-degree preferred orientation difference.

We fit a linear mixed-effects model and corrected p-values for multiple comparisons to study the main effect of orientation difference on SFC at 22 Hz (peak of the SFC). We found that orientation difference significantly affects SFC at 22 Hz (F (2.703, 500.9) = 16.65, p < 0.0001). Spike-LFP pairs with the same orientation preference had significantly higher SFC magnitudes than pairs with 30 degree (22 Hz, p = 0.0017,), 60 degree (p < 0.001) and 90 degree (p < 0.001) difference in preferred orientation. Pairs with 30 and 60 degree difference in preferred orientation had also significantly higher SFC magnitudes than pairs with 90 degree difference (p < 0.001 and p < 0.018, respectively).

To show the prevalence of the observed significant SFC effect, we also quantified the percentage of pairs showing significant SFC in each frequency band. The percentage of pairs showing significant SFC was higher for pairs with the same preferred orientation than pairs with different preferred orientations (**Fig. 2ci** and **2cii** for example and all monkeys, respectively). The percentage of significant SFC pairs peaked in the beta-band (**Fig. 2cii**, 72% at 22 Hz for all monkeys), and gradually decreased at higher frequencies, similar to the decrease in SFC magnitude across frequency.

Previous studies have noted SFC peaks in the gamma-band range in primary visual cortex (Siegel and König, 2003; Burns et al., 2010; Lashgari et al., 2012). These studies often calculate SFC from spikes and LFPs on the same electrode. Our peaks in the beta-band may be due to the minimum distance between electrodes (400um). To test this, we repeated the spike-LFP coherence analysis in all animals by taking spikes and the LFP from the same electrode. On a subset of sites (30/96, 31%) the peak of the SFC magnitude appeared in the gamma range (> 30 Hz), compared to ten sites when the LFP and spikes were obtained from neighboring electrodes (10/96 sites, 10 %). Therefore, the peak of SFC can be affected by the method by which it is computed.

### Distant orientation columns exhibit functional connectivity

Previous neuronal tracing studies have shown that distant orientation columns with the same orientation preference are preferentially connected (Gilbert and Wiesel, 1989; Malach et al., 1993; Bosking et al., 1997; Stettler et al., 2002). To test whether columns are functionally connected and whether functional connectivity across cortical distance changed as a function of orientation preference, SFC was calculated for all electrode pairs across the array (n = 11,225 pairs for four monkeys).

At each distance within each of the four preferred orientation difference groups, the percentage of pairs that showed significant SFC (*p* < 0.05, permutation test) was quantified. Although the percentage of pairs with significant SFC decreased as the distance between electrodes increased, pairs with the same orientation preference showed significantly higher SFC compared to the chance level (5%) in the beta band up to 2 mm (**Fig 3a**, example monkey, M1, **Fig 3b**, all monkeys.

**Fig. 3.**
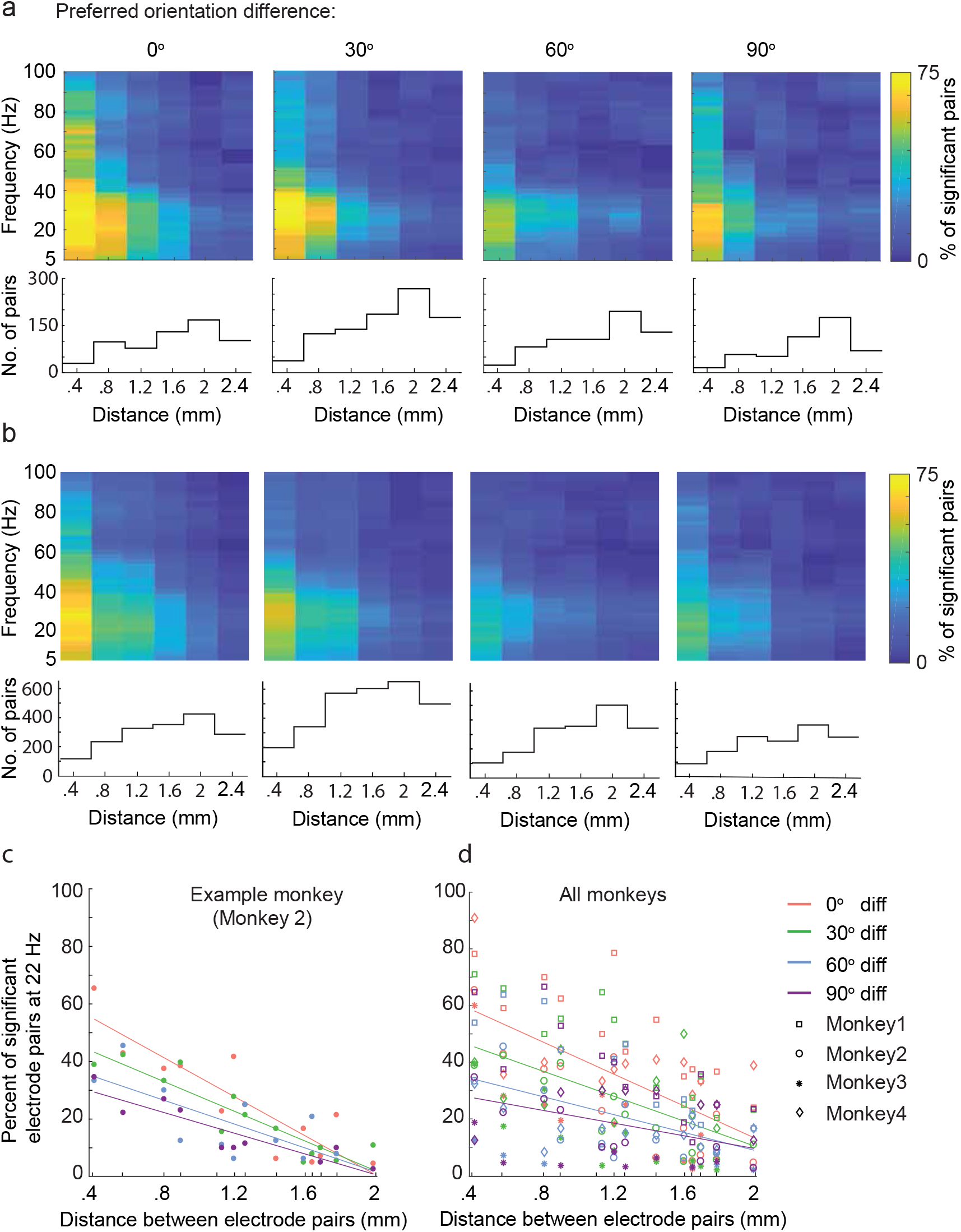
Spike-field coherency of adjacent neighboring sites. **a**, Percentage of pairs with significant spike-field coherence for binned distances (400 μm) and the total number of pairs in each binned distance for the example monkey and **b** averaged across four animals. The percentage of pairs with significant SFC decreases as the distance between pairs increase. **c**, Linear fit of the percentage of pairs with significant 22Hz spike-field coherence as a function of distance for the example monkey and **d**, for the population. The difference between orientation difference groups decreases as the distance between electrodes increases.

Since the distance between pairs of electrodes were not equal, we binned distances every 400 μm. At the shortest distance (400 μm), at least 50 per cent of pairs showed significant SFC in frequencies below 45 Hz where the maximum percentage of cells showing significant SFC was 75 percent at 22 Hz. As the distance between pairs increased, the magnitudes of SFC at 22 Hz dropped and fewer pairs showed significant SFC for outside of the beta-band. Low SFC across pairs separated by large distances was not due to the lower number of pairs in those groups, (**Fig. 3a,b**). Rather, the number of pairs in each group increased as the distance between electrodes increased.

To test the main effect of distance between recording pairs on the SFC peak, we fit a linear model on pairs with significant 22 Hz SFC across all electrode distances (Fig 3c for example monkey and 3d for all monkeys). Similar to SFC from neighboring electrodes, the percentage of distant electrode pairs showing a significant SFC at 22 Hz depended on the difference in preferred orientation of pairs. SFC decreased when LFP and spike sites had different preferred stimulus orientations. Furthermore, beyond a distance of 1.2 mm, differences in SFC between orientation groups was no longer significant (p = 0.7, Kruskal-Wallis test).

The observed SFC across cortical distance may be due to the general spread of current across cortex and not related functional processing within V1. In this way, decaying SFC across cortical distance in V1 should be similar to that of V1-V2 and V2-V2 pairings. To rule out this possibility, we simultaneously recorded from V1 and V2 sites in one animal and quantified the SFC from pairs within V1 or V2 and across V1-V2 at matched cortical distances. We fit a multiple comparison linear mixed-effects model to compare the main effect of cortical pairing (V1-V1, V1-V2 and V2-V2) and distance (400 to 1200 μm, binned every 400 μm) on SFC from pairs with the same stimulus orientation preference. The main effects were statistically significant for cortical pairing (F (2, 251) = 76.06, p < 0.001), distance (F (3.649, 509.0) = 24.80, p < 0.001), and there was a significant interaction (F (12, 837) = 20.04, p < 0.001). Tukey’s multiple comparison test showed that SFC at 22 Hz is significantly (p < 0.001) higher than that of V2 and V1-V2 pairs in 400 μm, 800 μm, 1200 μm bins. Therefore, the SFC we recorded across cortical distance was specific to processing within V1 where the decay in SFC across cortical distances is perhaps due to decrease in horizontal connections and not a general effect of cortical distance.

An earlier study (Jia et al., 2013) found a greater phase difference between V1 and V2 gamma rhythms compared to pairs of V1 sites. We repeated this analysis and measured the phase difference between V1 and V2 at 22 Hz. The circular mean phase difference for V1 (210 pairs) and V1-V2 (114) pairings were 28.1 ± 0.1 and 40.1 ± 0.3 degrees, respectively. This further suggests that our measures of functional connectivity are not simply due to cortical distance.

### Differences in spike-LFP coherence cannot be explained by firing rate alone

Differences in the spike-LFP coherency may be influenced by the spike rate. By definition, firing rates are higher at the preferred orientation. This means that a higher SFC magnitude in pairs with the same preferred orientation in Figure 2 could be due in part to a higher spike rate of pairs in this group.

To determine whether the differences we observed in SFC across orientation preference could be attributed to differences in spike rate, we selected pairs with a 0-degree and 90-degree difference with the same spike rate to compare their peak SFC together. To do so, pairs were grouped into two groups based on whether they had the same spike rate in their spike channels or LFP channels: spike-channel spike-rate-matched group including pairs with a 0-degree and 90-degree orientation difference that had the same spike rate in their spike channels grouped (**Fig. 4a**) and LFP-channel spike-rate matched group including pairs that had the same spike rate in their LFP channels (**Fig. 4b**). We correlated 22 Hz SFC from pairs with a 0-degree and 90-degree preferred orientation difference in both groups and found more points above the line of unity (dashed line), denoting higher SFC for pairs with a 0-degree preferred orientation difference compared to pairs with a 90-degree difference. We compared the SFC magnitudes of pairs with 0-degree and 90-degree preferred orientation difference (**Fig. 4c**) and found the highest SFC magnitudes when pairs had similar orientation preference in both spike-rate-matched groups (p = 0.0003, Spike channels, p = 5.4e-07, LFP channels). Therefore, the difference we observe in SFC magnitudes due to a difference in orientation preference cannot be attributed to differences in spike rate alone.

**Fig 4.**
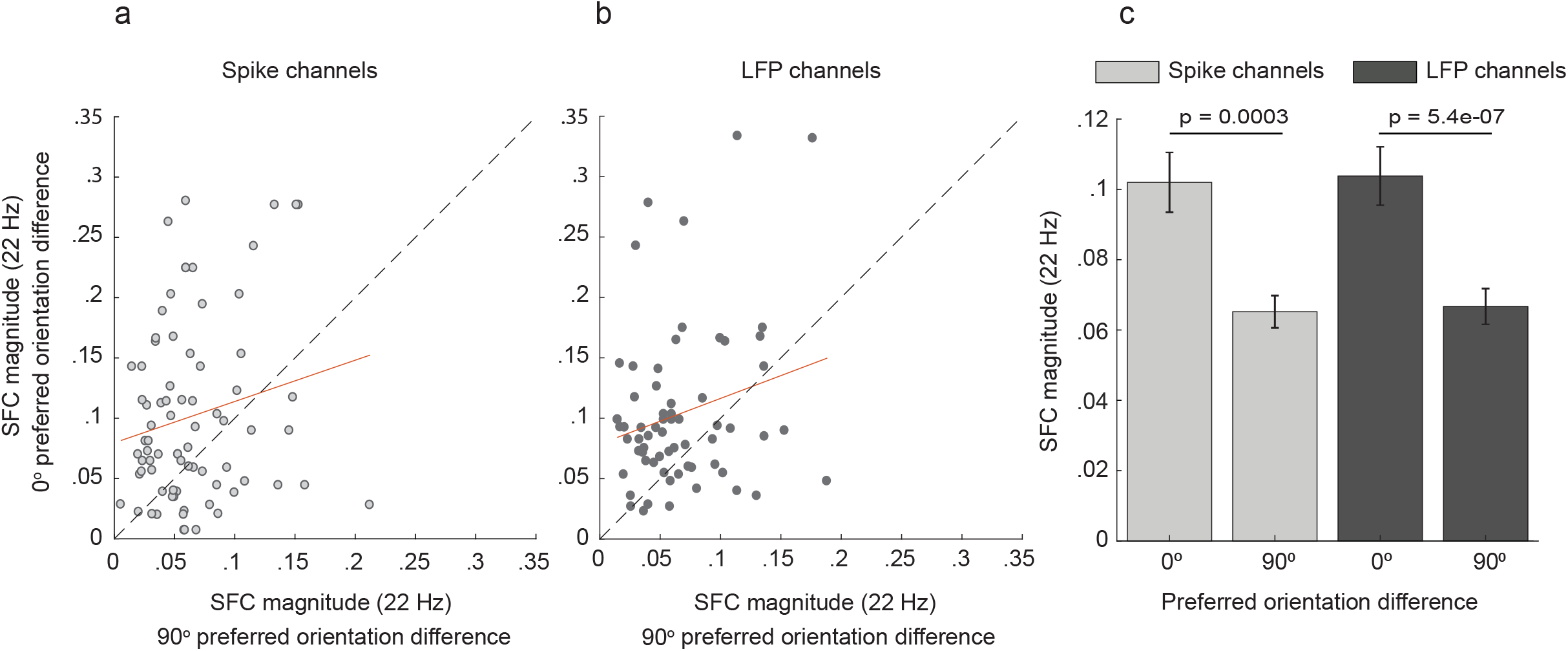
Spike-rate corrected spike-field coherence. **a,b**, Comparing SFC between pairs with 0- and 90-degree preferred orientation difference when spike rate was matched in their spike channels (n =) or the LFP channels (n =), respectively. **c**, In both control groups, pairs with 0-degrees preferred stimulus difference had significantly higher SFC magnitudes than pairs with a 90-degree difference. The dashed lines depict the line of unity.

### Pairs with higher SFC contain more decoded stimulus information

More information can be decoded from a neuron’s responses to a preferred stimulus compared to that of non-preferred stimuli (Kang, 2004). To determine whether the functional connectivity correlates with the amount of decoded information about the stimulus, we used an information-theoretic method to quantify how much information about the stimulus is shared between pairs of sites.

We grouped neurons’ spike count for all stimulus orientations (12 orientations, 900 trials) for each site and then measured the mutual information for each possible electrode pair. Next, we grouped the pairs based on their difference in preferred orientation difference. We found significantly higher mutual information from pairs with the same preferred orientation preference (0.61 ± 0.01 bits) than pairs with different preference (0.5 ± 0.015, 0.46 ± 0.012 and 0.42 ± 0.018 for 30 degree, 60 degree and 90 degree difference, respectively).

To connect these results with the results from the SFC analysis, we correlated spike-LFP pairs that showed significant SFC (22 Hz) with their mutual information values and fit a linear regression model to variables (**Fig. 5a_i_**, example monkey; **Fig 5a_ii_**, all monkeys. We found a significant correlation between SFC at 22 Hz and the mutual information across all preferred orientation differences (*r* = 0.047, *p* = 0.0002, Pearson’s correlation coefficient). Therefore, information about the stimulus orientation may be conveyed between orientation columns which can be detected by SFC in beta-band range.

**Fig 5.**
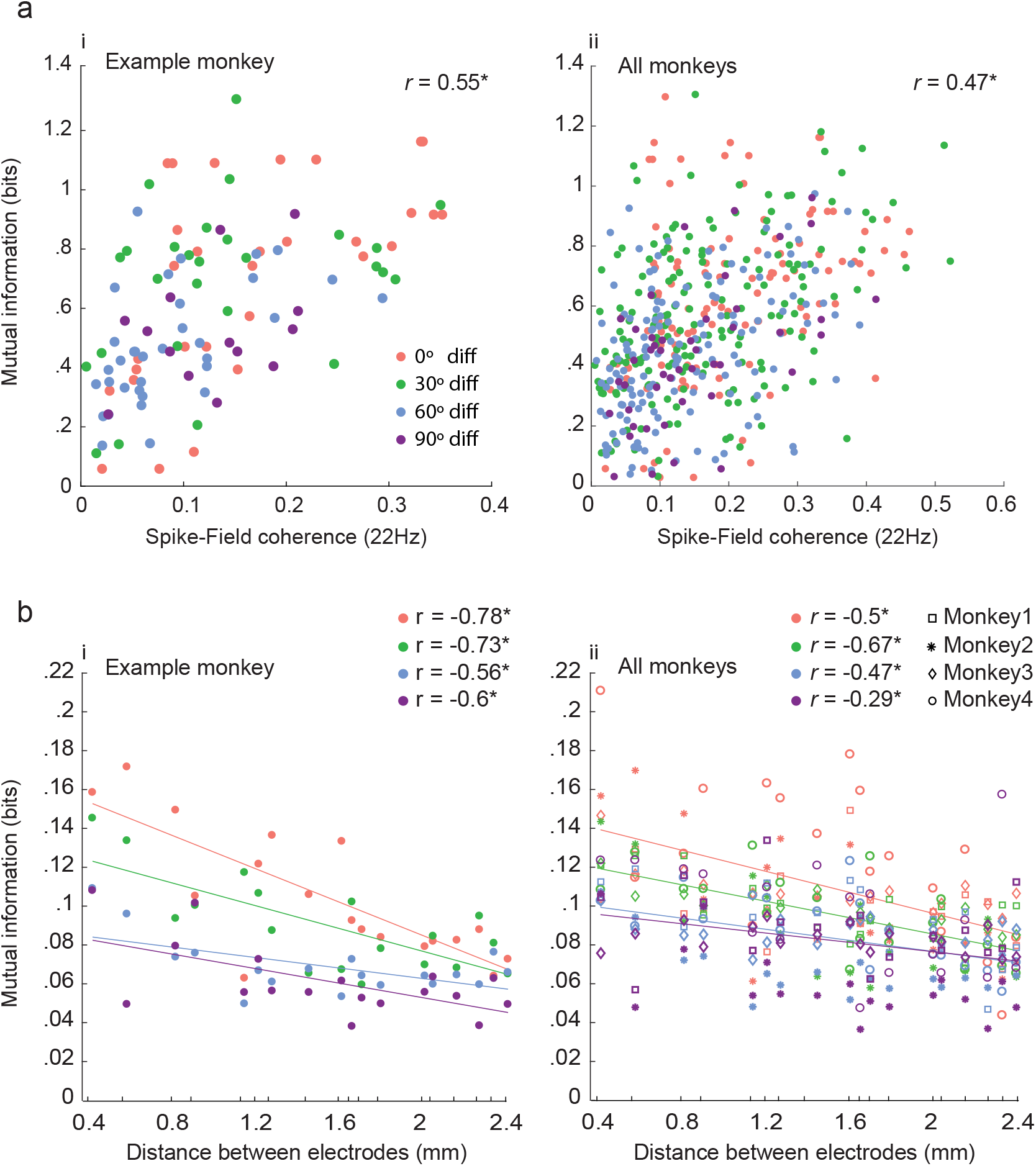
Information encoding in neighboring and distant sites. **a,b** Regression between mutual information and 22Hz SFC for neighboring sites. All orientation difference conditions are pooled together to measure regression value, however, conditions are color-coded for better observation. **c,d**, Mutual information for distant sites. Asterisks indicate significance (p < 0.05, F test).

Our results show that the amount of mutual information is significantly correlated with SFC at 22Hz. Given that increasing the distance between pairs results in a decrease in SFC, one may expect a similar decrease in stimulus information. To test this hypothesis, we measured how much mutual decodable information is affected by the distance between the electrodes. For each site, we grouped the neurons’ spike count for all stimulus orientations and then measured the mutual information for each possible electrode pair.

Based on the pairs’ stimulus preference difference, pairs were put into one of four groups according to their preferred orientation difference: 0, 30, 60 and 90-degree difference. We found that the distance between electrodes significantly affects the amount of encoded information between pairs of sites in all groups (p < 0.05). As the distance between the recordings sites increases, the mutual information decreases (**Fig. 5b_i_**, example monkey; **Fig 5b_ii_**, all monkeys).

The mutual information that we measure is akin to correlating the neurons tuning curves and significant correlation between SFC and MI denotes higher coherence between neurons with similar tuning curves. To make sure that the measured mutual information is mainly influenced by the similarity between pairings’ tuning cures and not by stimulus-induced variations in the spike rates, we calculated r_sc_ for each pair and measured Pearson correlation between r_sc_ and MI. We found significant positive correlation for sample monkey (r = 0.47, p = 0.01) and all monkeys (r = 0.5, p = 0.02)

## Discussion

We use a spike-LFP approach in this paper to study how neurons in orientation columns of V1 temporarily coordinate their activity patterns in response to a visual stimulus.

We quantified the SFC between neighboring (400 μm) and distant sites (up to 5.6 mm) in V1 and found that the magnitude of the SFC in pairs where both sites have the same preferred orientation preference is higher than that of pairs with different orientation preference. Higher SFC in spike-LFP pairs whose stimulus preferences were similar was not due to higher spike rates. Therefore, our results indicate that neurons in distant orientation columns can synchronize their activities where the level of synchronization depends on how much their stimulus preference is similar. To test whether synchronization between orientation columns can improve stimulus encoding, we used information-theoretic methods and found that more stimulus information can be decoded from spike-LFP pairs with the same preferred stimulus orientation than pairs with different preferred stimulus orientation where decoded information significantly correlated with SFC in the beta-band.

### Spike-LFP Coherence in V1

In the primary visual cortex, even neurons which are highly tuned for stimulus orientation have a broad tuning. Therefore, to bind the responses to the preferred stimulus together, neurons need to detect and distinguish responses to the preferred orientation from responses to the orientations close to the preferred orientation. Synchronization between neurons with the same stimulus responses has been proposed as a mechanism for selection of related responses. Our results support this notion by showing that when neurons whose receptive fields have a similar orientation preference are presented with a preferred visual stimulus, the coherence between their spike trains and the LFP increase. The increased SFC may indicate an increase in the strength of synaptic activity at one orientation column caused by spiking at another orientation column. Stronger synaptic connections may help the integration of synaptic inputs from neurons with the same orientation preference at postsynaptic neurons. It should be noted that peak SFC we report is lower (22 Hz) than that of previous studies which have shown an increase in the SFC in the gamma range (above 30 Hz) following visual stimulation of V1 in cats and monkeys (Siegel and König, 2003; Burns et al., 2010; Lashgari et al., 2012). This discrepancy can be explained by the difference in the SFC quantification in our experiments compared to the above studies where a single electrode was used to record spikes and LFPs simultaneously.

Although the LFP signal is dominated by summed synaptic activity in a cortical space surrounding the recording electrode (Mitzdorf, 1985; Buzsáki et al., 2012; Einevoll et al., 2013), there is a longstanding debate on the spurious effects of spikes on fast components of LFPs (Gold et al., 2006; Lepage et al., 2011). That means the magnitude of high-frequency SFC may be inflated if spikes and LFPs are recorded on the same electrode or from closely implanted electrodes. In our study, we quantified the SFC by taking spikes and the LFP from two adjacent electrodes which are at least 400 μm apart. When we repeated the spike-LFP coherence analysis by using spikes and the LFP from the same electrode in two monkeys, we found the peak of the SFC magnitude in the gamma range (> 30 Hz) in 31 % of sites as opposed to 10 % when SFC was quantified by taking spikes and the LFP from two adjacent electrodes. Based on our results, we suggest that SFC in gamma is, to some extent, due to the interference of spikes in the LFP and propose more caution while interpreting the gamma band SFC results. Our result also in line with the previous study (Lashgari et al., 2012) that reported two types of neurons being involved in SFC at high- and low-frequency bands which might be involved in different processing mechanisms (i.e. local vs long-range computation).

### Inter-columnar synchronization of neural responses

Although orientation columns with the same stimulus orientation preference are preferentially connected, lateral connections also connect sites with a wide range of preferred orientations (Kisvarday et al., 1989; Bosking et al., 1997; Stettler et al., 2002). Synchronization is a proposed way of enhancing communication between distant neuronal populations (Singer and Gray, 1995, 1995; Fries, 2005). Synchronization, however, needs to take place between the right neural populations, i.e. orientation columns with the similar orientation preference as synchronization between columns with different orientation preference may reduce the response binding efficacy in V1 (Singer, 1993; Singer and Gray, 1995). Central to this hypothesis is the premise that if spikes and the LFP of two neural clusters have the same rhythmicity, the maximum communication can be achieved when spikes sent by one neural cluster arrive when the other cluster is in its maximum sensitivity (Fries, 2005, 2015; Womelsdorf et al., 2007). Our data support this hypothesis by showing that the spike-LFP coherence depends on the orientation preference of recording pairs, highest SFC for pairs of sites with the same orientation preference and the least SFC for pairs with the maximum orientation difference, 90-degrees. Previous studies of neural synchronization have shown significant correlated oscillatory firing between remote orientation columns with the same orientation preference (Gray et al., 1989). Our results extend this observation in several important ways. Firstly, Grey et al. (1989) used single electrodes, which imposed a limited spatial resolution due to the number of electrodes they could implant simultaneously (4-6 electrodes). In our study, we used 96 equally distributed recording electrodes which significantly increases the spatial resolutions of recordings. Secondly, since the LFP measures accumulate synaptic activity of large pools of neurons (Mitzdorf, 1985; Okun et al., 2010), our measure – the spike-LFP coherence – is more sensitive for quantifying a phase synchronization than measures based on spike-spike synchronization. The higher sensitivity of the SFC method is particularly advantageous when studying long-range neuronal interactions. Finally, the synchronization between orientation columns reported in our study is supported by the results of an information theoretic (mutual information) and the interareal (V1-V2) functional connectivity methods.

The mutual information metric quantifies the amount of stimulus information obtained from one site through observing the information coded in another site (Quiroga and Panzeri, 2009). Therefore, the mutual information should not be affected by the distance between two sites if they encode the same stimulus and are totally independent. Our results showed decay in the mutual information when the distance between electrodes increases. This can potentially be explained by the horizontal connections between columns. Each orientation column receives extensive inputs from distant orientation columns with the same preferred orientation and some inputs from neighboring columns with different orientation preference, however, the horizontal connection probability decrease when the distance increases (Bosking et al., 1997; Stettler et al., 2002). Therefore, pairs of columns with a large distance may receive more inputs from their non-iso-oriented columns arriving from close neighboring columns which eventually reduce the mutual coding of visual stimuli.

Our results show that synchronization between V1 neuron’s spikes and the LFP from a neural cluster up to 2mm apart happens when both neural clusters have the same stimulus orientation preference. These results can be explained by the physiology of horizontal connections which preferentially link remote clusters of neurons with similar orientation preference (Burkhalter, 1989; Gilbert and Wiesel, 1989; Kisvarday et al., 1989; Schwarz and Bolz, 1991, 1991; Kisvárday and Eysel, 1992; Malach et al., 1993, 1993; Bosking et al., 1997). The synchronization between columns as separate as 2mm apart may help the flow of information to the downstream as several studies have shown that temporarily correlated activity of large pools of neurons can help the control of the flow of information across cortical areas (Salinas and Sejnowski, 2001; Rubino et al., 2006).

Orientation columns in V1 are also connected through local/short-range horizontal connections which are shorter than 500 um in macaques (Stettler et al., 2002) and are less specific as they connect neurons with different orientation preference. This means the SFC magnitude up to 500 μm in the macaque’s brain should not be affected by the stimulus preference of orientation column pairs which seems to be in contrast with our results. However, it should be noted that functional columns in marmoset V1 are much smaller than the macaque brain, so as the horizontal connections (Roe et al., 2005; McLoughlin and Schiessl, 2006). Therefore, the neighboring columns in our study with 400 μm separation are most likely connected mainly through long-range horizontal connections.

Finally, we investigated synchronization between orientation columns with similar stimulus orientation preference within V1, within V2 and between V1 and V2. We found a larger SFC at 22 Hz between V1-V1 pairings compared to V1-V2 and V2-V2 pairings. These results are in line with the notion that corticocortical connections within V1 are higher than within V2 and between V1 and V2 (Jia et al., 2013). Therefore, higher SFC magnitudes for V1 pairings are expected compared to other pairings. The larger SFC between pairs of V1 neurons also implies that our results are related to functional connectivity between orientation columns and not merely an effect of signals propagating across the cortical surface. Our results, however, are different from previous interareal studies showing the strongest synchronization between V1 and V2 cells whose their receptive fields overlap and the orientation preferences are similar (Nowak et al., 1999; Jia et al., 2013). By measuring cross-correlation between cells with overlapped receptive fields, the purpose of the above studies was to show how synchronization between areas can alter the efficacy of drive from V1 to V2. We did not match the receptive fields of pairs as the purpose of our study was to understand how distributed groups of neurons within V1 synchronization their activities following simultaneously being visually stimulated.

## Acknowledgements

We acknowledge the Australian Research Council (DE180100344, DP200100179) and the National Health and Medical Research Council (APP1185442, APP1120667) for financial support. We also thank Janssen-Cilag for the donation of sufentanil citrate which made our experiments possible.

